# β-catenin localization in the ctenophore *Mnemiopsis leidyi* suggests an ancestral role in cell adhesion and nuclear function

**DOI:** 10.1101/2024.08.29.610370

**Authors:** Brian M. Walters, Lucas J. Guttieres, Mayline Goëb, Stanley J. Marjenberg, Mark Q. Martindale, Athula H. Wikramanayake

## Abstract

The emergence of multicellularity in animals marks a pivotal evolutionary event which was likely enabled by molecular innovations in the way cells adhere and communicate with one another. β-catenin is significant to this transition due to its dual role as both a structural component in the cadherin-catenin complex and as a transcriptional coactivator involved in the Wnt/β-catenin signaling pathway. However, our knowledge of how this protein functions in ctenophores, one of the earliest diverging metazoans, is limited. To study β-catenin function in the ctenophore *Mnemiopsis leidyi*, we generated affinity-purified polyclonal antibodies targeting *Ml*β-catenin. We then used this tool to observe β-catenin protein localization in developing *Mnemiopsis* embryos. In this article we provide evidence of consistent β-catenin protein enrichment at cell-cell interfaces in *Mnemiopsis* embryos, suggesting that *Ml*β-catenin had an ancestral role in cell adhesion. Additionally, we found β-catenin enrichment in some nuclei, particularly restricted to the oral pole around the time of gastrulation, suggesting *Ml*β-catenin may have a nuclear function in *Mnemiopsis* embryos. The *Ml*β-catenin affinity-purified antibodies now provide us with a powerful reagent to study the ancestral functions of β-catenin in cell adhesion and transcriptional regulation.

## 1. Introduction

The origin of multicellularity in animals represents a major evolutionary transition, marking the divergence from single-celled ancestors to complex organisms composed of multiple, specialized cell types. Understanding the mechanisms that evolved to enable this transition will provide insights into the fundamental principles that underpin animal evolution and the diversity of life forms we see today. Critical innovations such as cell-cell adhesion, cell-extracellular matrix interactions, cell-cell communication, and cell fate specification are hypothesized to have facilitated cellular cooperation and the partitioning of roles in the cells of the metazoan last common ancestor.^1,2^ The β-catenin protein is a multifunctional molecule central to many of these cellular processes and thus stands out as a key protein for understanding this transition.^3,4^

β-catenin is a structurally and functionally conserved protein found to be important for developmental processes and maintaining tissue homeostasis across many metazoan lineages. It is well established that β-catenin plays two important roles: acting as a structural component of the cadherin-catenin complex (CCC) that maintains cell-cell adhesion through adherens junctions (AJs) and as a transcriptional coactivator in the Wnt/β-catenin signaling pathway.^4^ Because of its dual purpose, β-catenin has been identified as a key component involved in cell adhesion, cell fate specification, neurogenesis, spindle orientation, cell migration, cell polarity, and maintenance of stem cells.^4^

In its role in cell-cell adhesion, β-catenin interacts with cadherin, a calcium-dependent transmembrane glycoprotein that links neighboring cells together by forming dimers in the extracellular space.^5^ Cadherins alone are not sufficient in the establishment of a stable AJ.^6^ The intracellular domain of classical cadherins includes binding sites for two catenins: p120 and β-catenin which are essential for stabilizing this complex at the plasma membrane (Fig. 1).^7–10^ In the absence of binding, members of the CCC are degraded through multiple mechanisms, destabilizing the AJs and leading to a loss of adhesion between cells.^11–13^ This interaction plays a pivotal role in maintaining tissue integrity and organization in multicellular organisms.^14^ Thus, β-catenin is essential in cell-cell adhesion through the formation and maintenance of AJs by interacting with cadherins.

**Fig. 1.**
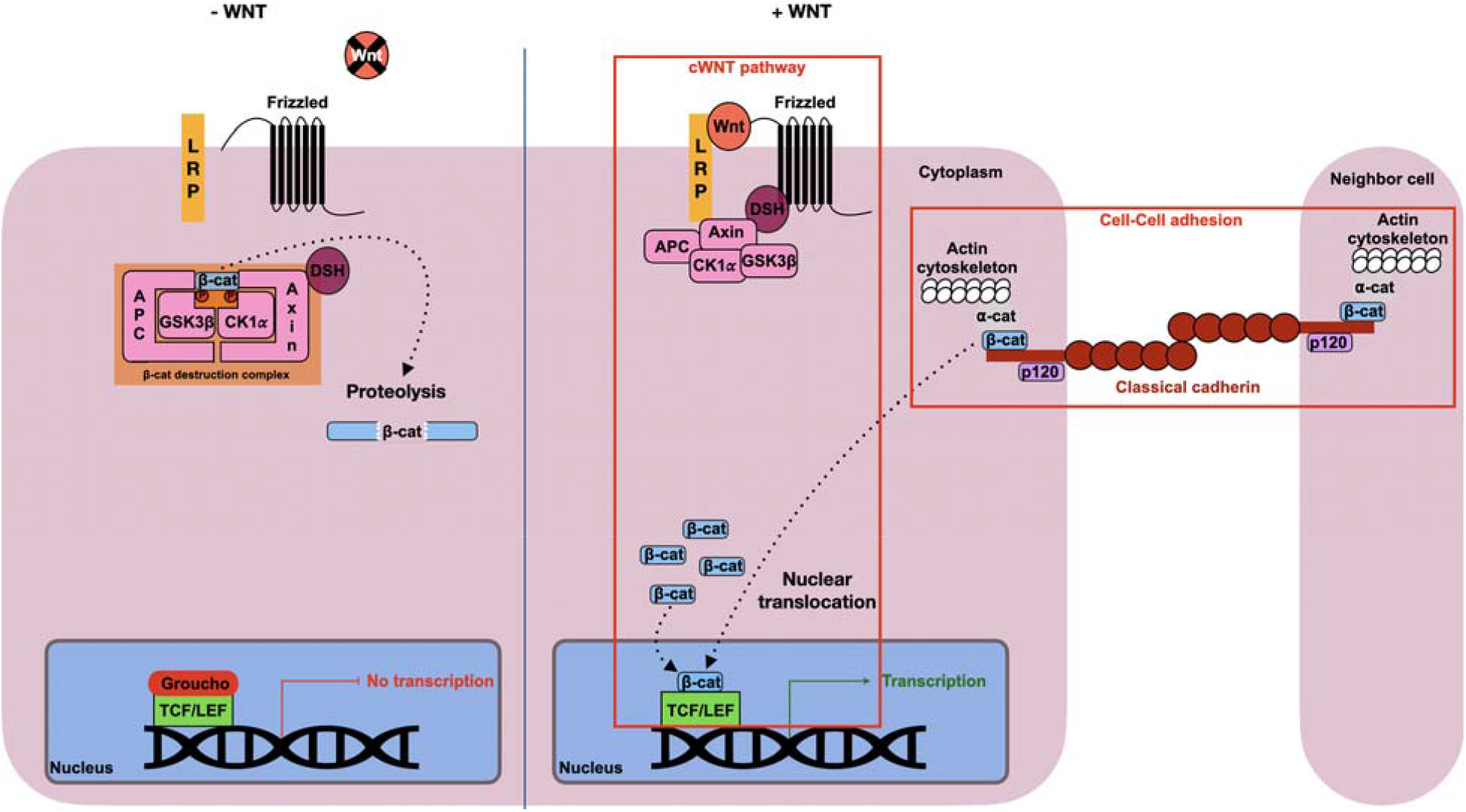
The dual role of β-catenin in cell adhesion and transcriptional activation. LRP, Low-density lipoprotein Receptor-related Protein; APC, Adenomatous Polyposis Coli; CK *α*, Casein Kinase 1*α*; GSK-3β, Glycogen Synthase Kinase 3β; TCF, T-Cell Factor; LEF, Lymphoid Enhancer Factor; Dsh, Dishevelled; β-cat, β-catenin; *α*-cat, *α*-catenin.

On the cytoplasmic side of the membrane, β-catenin has a role in recruiting ⍰-catenin (⍰-cat) to the CCC, enabling interaction between the plasma membrane and the actin cytoskeleton. This interaction is necessary for cell adhesion and provides the mechanical force that maintains tissue integrity.^15^ In turn, this mechanical force exerted on the CCC is at the heart of the mechano-transduction process, which is characterized by the translation of a mechanical stress into a biochemical message. This mechanically regulated cadherin/β-catenin signaling pathway is notably involved in the specification of the anterior endoderm in *D. melanogaste*r and in the activation of key genes in mesoderm specification by β-catenin in zebrafish and Drosophila.^16–18^ Furthermore, there is evidence that selective nuclearization of CCC associated β-catenin can be induced through tension-relaxation events, leading to β-catenin facilitated transcription.^19,20^

Furthermore, β-catenin plays an important role in the Wnt/β-catenin signaling pathway, which can be maintained in two states: ‘off’ and ‘on’ (Fig. 1).^21^ In the absence of the Wnt ligand, the pathway remains ‘off’. Here a multi-protein complex, known as the destruction complex, regulates cytoplasmic β-catenin stability. Within this complex, Axin and Adenomatous Polyposis Coli (APC) serve as recruitment and scaffolding proteins while Casein Kinase 1 alpha (CK1*α*) and Glycogen Synthase Kinase-3β (GSK-3β) sequentially phosphorylate β-catenin, marking it for degradation through the proteasome pathway.^22,23^ On the other hand, when the Wnt ligand is present the pathway shifts to the ‘on’ state. The Wnt ligand binds to the Frizzled (Fzd) and Low-density lipoprotein Receptor-related Protein (LRP5/6) transmembrane co-receptors which then each respectively recruit Dishevelled (Dsh) and Axin to the cell membrane. Receptor inhibition of GSK-3β and Axin phosphorylation interfere with the destruction complex’s enzymatic activity, allowing cytoplasmic β-catenin to accumulate. At a high enough concentration, β-catenin is able to translocate to the nucleus where it binds to T-Cell Factor/Lymphoid Enhancer Factor (TCF/LEF) transcription factors and acts as a transcriptional coactivator. The ultimate outcome of β-catenin nuclearization during the ‘on’ state of the pathway is the transcription of downstream target gene expression. Mutations and altered expression patterns of this pathway have been linked to human diseases including cancer.^24,25^

During the early development of many bilaterian animals, Wnt/β-catenin signaling is involved in the establishment of the primary embryonic axis, known as the animal-vegetal (AV) axis.^26,27,28^ The AV axis is significant because it orients the embryo for future developmental processes, such as germ layer specification.^27,28^ During normal development of some bilaterians, nuclearization of β-catenin at the vegetal pole specifies the vegetal blastomeres to give rise to endomesoderm, a bipotential germ layer that will later segregate into endoderm and mesoderm.^28,29^ Development can be manipulated by inhibiting the nuclearization of β-catenin in the vegetal micromeres and macromeres in sea urchin and nemertean embryos-disrupting endomesoderm specification and causing ectopic expression of anterior neuroectoderm markers throughout the embryo-resulting in an “anteriorized” phenotype.^29–33^ Conversely, ectopically activating the Wnt/β-catenin pathway in the anterior mesomeres breaks the established polarity causing cells typically fated to become ectoderm to become respecified as endomesoderm, giving the embryo a “posteriorized” phenotype.^30,34^ However, while this pattern is widely observed in bilaterians, not all metazoans specify endomesoderm at the vegetal pole.^26,28^ Cnidarians, the closest outgroup to the bilaterians, establish a primary oral-aboral (OA) axis which is patterned by Wnt/β-catenin signaling activation at the oral pole.^35,36^ Other studies in a more basal metazoan, the ctenophore, has shown that endomesoderm specification also takes place at the oral pole.^37,38^ However, it is not entirely clear if this specification sets up the OA axis in these animals.

Most of our understanding of β-catenin’s functional role in metazoan biology comes from bilaterian and cnidarian models.^4,39–41^ While these studies are important for establishing an understanding of these highly conserved processes for β-catenin, investigating earlier diverging taxa will provide valuable insights into the evolutionary origins of these mechanisms. For instance, the role of β-catenin in either a cell signaling or cell-cell adhesion context in ctenophores remains poorly characterized. Several lines of evidence now indicate that the ctenophores, or comb jellies, were the earliest emerging metazoan clade.^42– 45^ Ctenophores have a small genome (for example, the *Mnemiopsis leidyi* genome is roughly 150 megabases) with a reduced molecular repertoire relative to other metazoans, which simplifies the identification of conserved proteins and interactions necessary for molecular function. For example, initial *in silico* analysis suggests that ctenophore proteins such as Cadherin and Axin lack the domain responsible for binding to β-catenin,^36,46^ and there is no evidence for a Wnt/Planar Cell Polarity (PCP) signaling pathway.^47,48^ This raises questions about what type of information will emerge from studying ancestral protein functions in ctenophores. It could either narrow down essential domains required for structural interactions or uncover alternative methods employed for accomplishing developmental and homeostatic processes. Moreover, studying key proteins in ctenophores can enhance our understanding of their evolutionary origins and shed light on how these essential biological processes evolved across Metazoa. Ctenophores, like *M. leidyi*, occupy a critical phylogenetic position, offering potentially unique insights into the molecular transition from unicellularity to multicellularity in the animal lineage.

Therefore, developing tools to study β-catenin and other critical molecules in ctenophores will be instrumental in uncovering these insights. Here we report on the successful generation of affinity-purified rabbit polyclonal antibodies targeting *M. leidyi* β-catenin protein and use it to carry out localization studies to determine the subcellular distribution of the protein during early *M. leidyi* development. Using these antibodies we were able to observe β-catenin protein localize to the cell-cell interface in the earliest stages of development and in some subsets of nuclei of developing *M. leidyi* embryos. This reagent provides a useful tool for future studies on the potential function of β-catenin in cell-cell adhesion and transcriptional activation in this member of an early emerging metazoan taxon.

## 2. Materials and Methods

### 2.1 Structural analysis of *Ml*β-catenin protein domains

The protein domains of *Ml*β-catenin (*Ml*β-cat, ML073715a) have been predicted and checked with Interpro 100.0 (https://www.ebi.ac.uk/interpro/)^49^ and SMART 9.0 (http://smart.embl-heidelberg.de/)^50^. Tertiary structure was predicted using AlphaFold v2.3.1 on the COSMIC^2^ platform (https://cosmic2.sdsc.edu/)^51^.

### 2.2 Maintenance and spawning

The *M. leidyi* adults were caught in a marina at Flagler Beach (Florida, United States, 29°30’60.0”N 81°08’47.1”W) and were maintained in 40L kreisels with constant seawater flow, temperature, and salinity at the Whitney Laboratory for Marine Bioscience of the University of Florida (Saint Augustine, Florida, USA). Care and maintenance of adults including spawning, fertilization, and embryo culturing have been carried out as previously described,^52^ however the salinity the adult animals were kept in was adjusted to ⅔ strength filtered sea water (FSW, 25 parts per thousand) to improve spawning efficiency. Spawning was induced by incubating the adults for 3 hours in the dark at RT and then moving them to glass bowls approximately one hour before spawning to collect embryos. Embryos were kept in glass dishes in ⅔ FSW until the desired stage.

### 2.3 Molecular cloning of *Ml*β-cat

The total RNA was extracted from mixed embryonic stages using Trizol (Sigma, cat.# T9424). A partial length *Ml*β-cat coding sequence was amplified from cDNA using the Advantage RT for PCR kit (Clontech, cat #639506) using the following primers:

*Ml*β-cat pGEX forward: ATAGGATCCATGGAAACGCCAGTATAT

*Ml*β-cat pGEX reverse: TATAAGAATTCTCAGGTCGTTGTGACGGG

The amplified cDNA was then ligated into a pGEX-6P-1 vector for cloning and inducing a GST-tagged *Ml*β-cat protein. The first 200 amino acids of *Ml*β-cat were selected to be used as the antigen for the polyclonal antibodies due to its predicted high immunogenicity. BLAST search against the *M. leidyi* Protein Model 2.2 (*Mnemiopsis* Genome Project Portal, https://research.nhgri.nih.gov/mnemiopsis/) using the antigen sequence only recovered the β-catenin sequence at a significant E-value.

### 2.4 Antigen preparation and antibody generation

BL21(DE3) competent *E. coli* (New England Biolabs cat.# C2530H) were transformed with the *Ml*β-cat pGEX-6P-1 construct. Five milliliters of inoculated terrific broth (TB) medium were incubated overnight (12-18 hours) in a shaking incubator at 37°C at 200 rpm. Overnight cultures were used to inoculate 500 mLs of TB medium which were then incubated until they reached an OD_600_ between 0.2-0.3, at which point the bacteria were induced by adding 0.5 mM isopropyl β-d-1-thiogalactopyranoside (IPTG). Cultures were allowed to incubate for an additional 3 hours before bacteria were collected by centrifuging at 5000g for 15 minutes at 4°C. The induced GST-tagged *Ml*β-cat protein was affinity purified on a glutathione sepharose beads matrix as described.^53^ Approximately 1.5 mg of Mlβ-cat protein collected via PreScission Protease (GenScript cat.# Z02799) digest and 6 mg of *Ml*β-cat-GST protein collected via 10 mM glutathione whole protein elution were sent to Labcorp Early Development Laboratories Inc. (Denver, PA). There, the *Ml*β-cat N-terminal polypeptide was used as the antigen for single rabbit polyclonal antibody generation and the *Ml*β-cat-GST protein was used for affinity purification of the antibody from final bleed serum.

### 2.5 Western blot analysis

*M. leidyi* embryo protein samples were collected by centrifuging mixed stage embryos (between 8 and 12 hours post fertilization, hpf) in an Eppendorf tube to pellet, removing as much sea water as possible, and adding minimum volume (between 30-50 µL) of 1X SDS gel-loading buffer (50 mM Tris-Cl pH 6.8, 100 mM dithiothreitol, 2% sodium dodecyl sulfate, 0.1% bromophenol blue, 10% glycerol) to fully lyse the pellet. *M. leidyi* adult protein samples were collected by dissecting the lobes off, using a homogenizer to dissociate cells, centrifuging to pellet, removing as much sea water as possible, and adding minimum volume (between 100-500 µL) of 1X SDS gel-loading buffer to fully lyse the sample. Lysates were boiled for 5 minutes and 30 µL was then separated on 10% SDS-PAGE gels, transferred onto 0.45µm nitrocellulose membranes (Bio-Rad Laboratories, Inc. cat. # 1620115) and blocked overnight in a 5% milk powder 1X tris buffered saline solution. The immunoblot was probed with the rabbit anti-*Ml*β-cat (1:500). Blots were developed using a IRDye 800 (1:10,000) donkey anti-rabbit secondary antibody (LI-COR, Inc. cat.# 926-32213) and imaged using a LI-COR Odyssey Infrared Imaging System.

### 2.6 Fixing and immunofluorescence analysis of embryos

The vitelline membranes of *M. leidyi* embryos were mechanically removed using forceps before fixation. Embryos were kept in plastic dishes treated with 3% Bovine Serum Albumin (BSA) in ⅔ FSW to prevent them from sticking to the dish. Embryos of different developmental stages were fixed (4% paraformaldehyde, 0.2 % glutaraldehyde in 1X phosphate buffered saline) 5 minutes at RT, before being fixed in a second fixative solution (4% PFA in 1X phosphate buffered saline) 10 minutes at RT. Embryos were washed three times with 1X PBS then permeabilized in 0.2% Triton in PBS. Embryos were rinsed several times in 1X PBT (0.1% BSA, 0.2% Triton in 1X phosphate buffered saline) before the blocking step in 5% Normal Goat Serum (NGS in 1X PBT) for 1 hour at RT on a rocker. The blocking solution was then replaced by the primary antibody (anti-*Ml*β-cat; 1:500 in blocking reagent) and incubated overnight at 4° with gentle rocking. Fixed embryos were washed several times with 1X PBT. A goat anti-rabbit IgG AlexaFluor 647 (Invitrogen,cat.# A-21245) secondary antibody was diluted in 5% NGS (1:250) and embryos were incubated in the dark at 4° overnight on a rocker. Negative controls were incubated with the secondary antibody only. Samples were washed with 1X PBT for a total period of 2 hours. Embryos were finally stained with DAPI (1:1,000 in PBT; Invitrogen, Inc. Cat. #D1306) to allow nuclear visualization. Stained samples were rinsed 2 times with 1X phosphate buffered saline before mounting in the same solution. The embryos were imaged under a Zeiss 710 scanning confocal microscope. All the images were processed through ImageJ (z-stack and maximal intensity projection). More than 30 embryos were examined at each embryonic stage.

## 3. Results

### 3.1. β-catenin protein structure in M. leidyi

In most metazoans, the core of the β-catenin protein is composed of 12 Armadillo repeat domains (Arm), essential for the interaction with β-catenin binding partners for both Wnt/β-catenin signaling and in AJs.^41,54^ Those binding sites are overlapping, meaning that β-catenin can either bind to cadherin or TCF/LEF, but cannot simultaneously accomplish both functions.^8^ The *M. leidyi* genome encodes one single β*-catenin* gene of 2,688 bps coding for 895 amino acids.^47^ Protein domain prediction softwares were only able to identify 9 out of 12 individual Arm repeats conserved in *Ml*β-cat, but also highlighted a larger Arm repeat superfamily that extended past the final recognized domain (Fig. 2A). Upon further investigation into the tertiary structure, 3 additional complete triple alpha helices were identified with relatively strong per-residue local confidence (Fig. 2B and C). Arm domain 7 was predicted with low confidence likely due to only 2 out of the 3 alpha helices being conserved in predicted 3D structure. However, a third, short alpha helix is present immediately before the domain but truncated by a low confidence value region of disorder. Furthermore, the alignment of *Ml*β-cat with structurally characterized orthologs in other animals revealed the conservation of the 2 lysine residues (K312 and K435 in mouse epithelial cadherin) in *Ml*β-cat, crucial for the interaction between a classical cadherin and β-catenin.^9^ Moreover, the phosphorylation sites required by β-catenin destruction complex components, part of the Wnt/β-catenin pathway, are also conserved (Serine 112, Serine 116, and Threonine 120 for GSK3β ; Serine 124 for CK1*α*). Additionally, 9 out of the 14 essential binding residues of the ⍰ -catenin binding motif are conserved.^46^

**Fig. 2.**
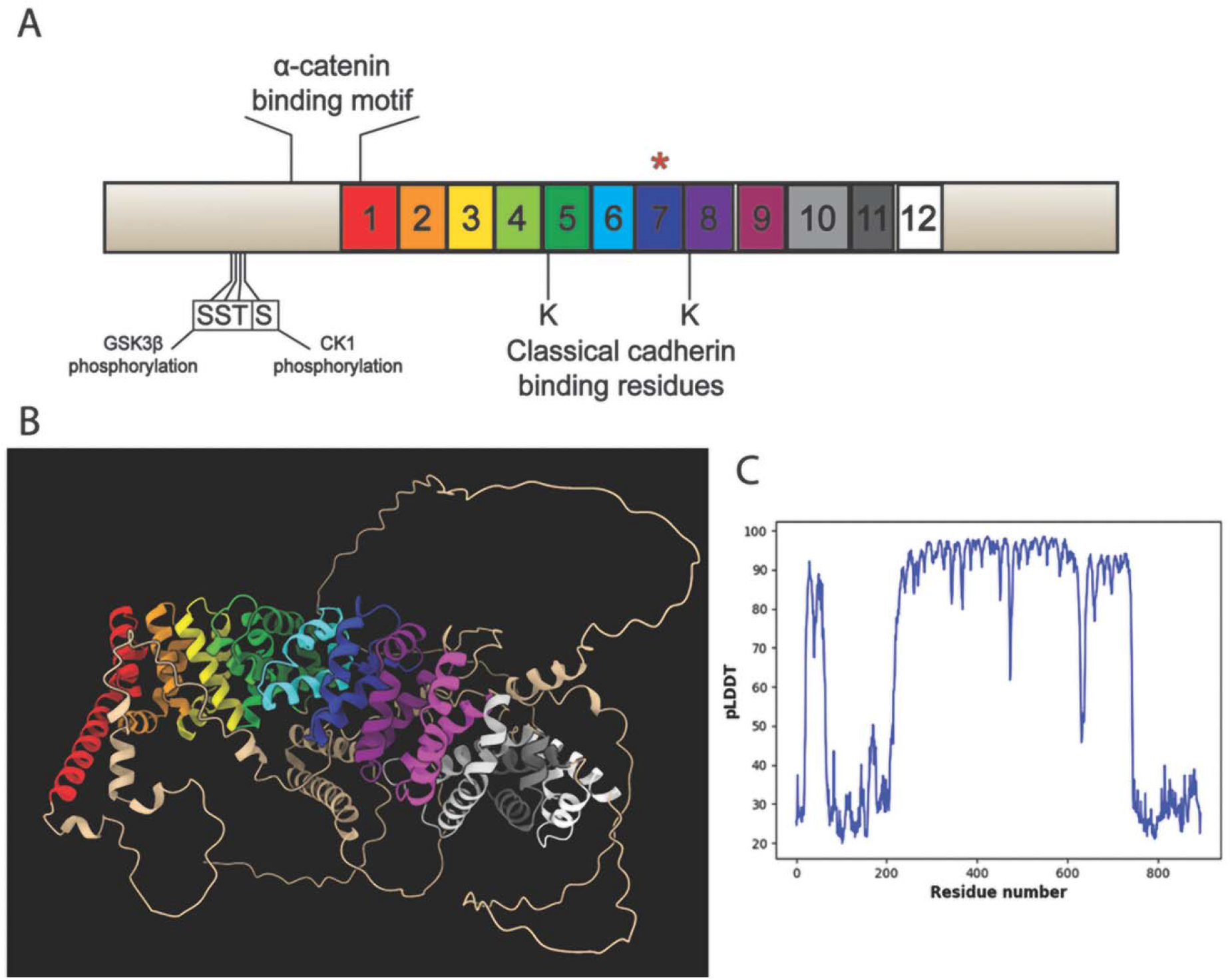
The predicted β-catenin architecture in *M. leidyi*. **(A)** *Ml*β-cat protein contains 11 of the 12

Arm domains identified with high confidence flanked by a N-Terminal and a C-terminal region. Asterisk above domain 7 indicates predicted partial (2 out of 3 alpha helices) domain conservation. The amino acids necessary for the interaction with GSK3β and CK1*α* are conserved, as well as the *α*-cat binding site. The two crucial lysines (K), essential for the interaction with a classical cadherin, are also conserved. **(B)** Predicted tertiary structure of *Ml*β-cat protein with Arm domains denoted by color. Red through pink domains were identified through domain prediction software; silver, gray, and white domains fall within the Arm repeat superfamily but were identified manually. The blue Arm domain (domain 7) was identified with low confidence likely due to the missing alpha helix. **(C)** pLDDT plot for predicted structure. Dip in confidence values for residues between 472-478 corresponds with a disordered region that truncates an alpha helix in Arm domain 7.

### 3.2 Affinity-purified rabbit polyclonal antibodies generated to Mlβ-catenin protein specifically targets endogenous Mlβ-catenin

To confirm that the affinity-purified rabbit polyclonal antibodies generated were specifically targeting *Ml*β-catenin, whole cell lysates from both embryo and adult *M. leidyi* were collected and used for Western blot analysis. Only a single band was detected around the expected size (∼100 kDa) for both samples (Fig. 3A). The discrepancy in size could be explained by the posttranslational modification of *Ml*β-catenin since the protein is known to be phosphorylated in other species,^23^ which is known to impact the outcome of protein size on Western blots. Next, fixed whole embryos from multiple stages were immunostained using the *Ml*β-cat antibody and nuclei were labeled with DAPI. Fixed embryos incubated with the secondary antibody alone were used as a negative control. Samples imaged using scanning confocal microscopy demonstrated robust nuclear staining by DAPI, but showed fluorescence only when embryos were incubated with the primary *Ml*β-cat antibody (Fig. 3B).

**Fig. 3.**
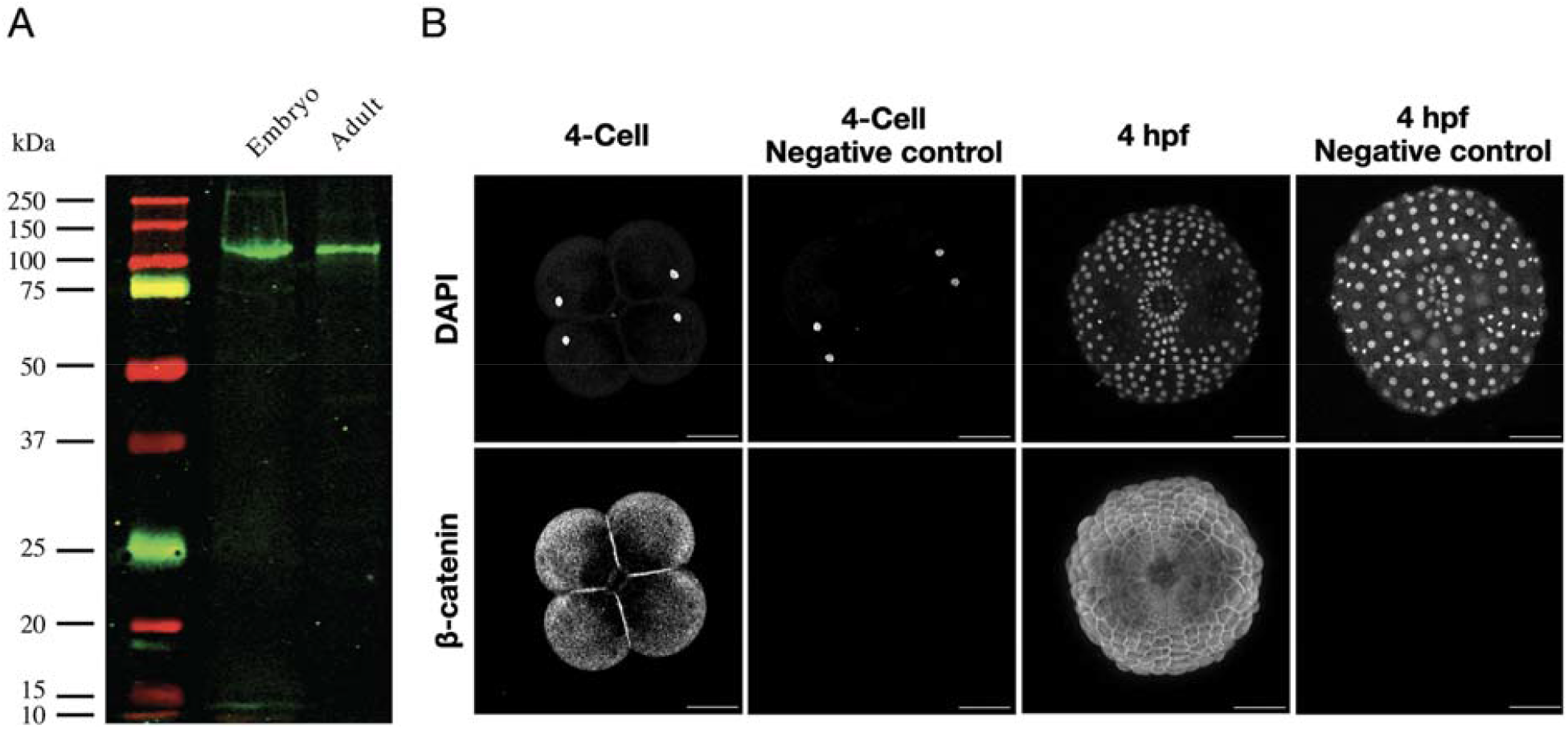
*Ml*β-cat antibody specifically recognizes endogenous protein. **(A)** Western blot analysis of *M. leidyi* cell lysates using affinity-purified anti-*Ml*β-cat antibodies. The specificity of the antibody was tested against whole-cell lysates from both *M. leidyi* embryos and adults. A single band slightly larger than the expected size (∼100kDa) was detected for both samples. **(B)** Immunostaining of *M. leidyi* embryos using affinity-purified anti-*Ml*β-cat polyclonal antibodies. The top panels show nuclei stained by DAPI; the bottom panels show results of *Ml*β-cat staining with and without primary antibody (negative control). Images are maximal projection. Scale bar, 50 µM.

### 3.3 Mlβ-cat localizes at cell-cell contacts

β-catenin is known to play a structural role by its importance in cell-cell adhesion by its association with cadherins at the plasma membrane. To determine if *Ml*β-cat was putatively involved in cell-cell adhesion, we stained *M. leidyi* embryos at different cell-stages from fertilized egg to 8 hpf. We were not able to detect any membrane staining of *Ml*β-cat at the 1-cell stage, however a cortical localization of *Ml*β-cat at cell-cell contacts at the 2-cell stage was visible. This localization continued from the 2-cell stage through the late stages of the development in *M. leidyi* (Fig. 4). This localization pattern was only present at cell-cell contacts, while the region of cell membranes that lacked contact displayed a diffuse level of *Ml*β-cat staining that more closely resembled the cytoplasm.

**Fig. 4.**
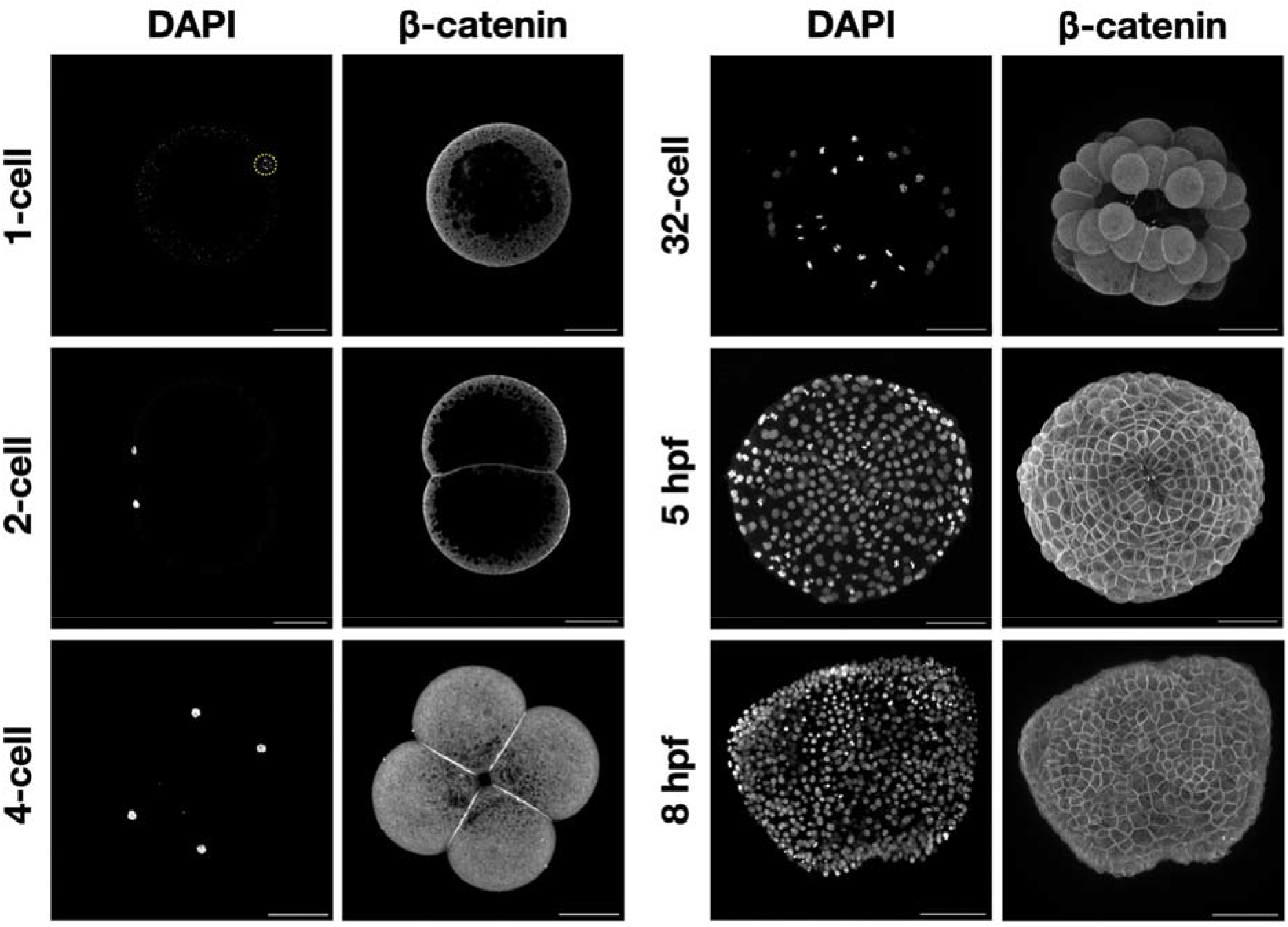
*Ml*β-catenin localizes at cell-cell contacts during *M. leidyi* embryonic development. Immunostaining of *M. leidyi* embryos at different stages using affinity-purified β-cat antibodies. The left column panels exhibit nuclei stained by DAPI; the right column panels exhibit results of *Ml*β-cat antibody staining. Images for 1 and 2-cell stages are z-stack projected images. All other images are maximal projection. Note that enriched localization of β-catenin only occurs at cell-cell contacts. Scale bar, 50 µM.

### 3.4 Mlβ-cat translocates into the nucleus during early embryogenesis

β-catenin function in cell signaling is associated with its accumulation into the nucleus to act as a transcriptional cofactor with TCF/LEF. We therefore looked for nuclear staining using the *Ml*β-cat antibody. We first noticed that *Ml*β-cat was uniformly expressed in the cytoplasm at 1-cell stage, indicating that this protein is likely a maternal component in *M. leidyi*. However, we did not consistently detect any clear nuclear signal of β-catenin from 1-cell to 32-cell (Fig. 5). The first *Ml*β-cat nuclear translocation event that we observed was at the 60-cell stage in some, but not all, 3E and 2M\ macromeres as well as some micromeres (Fig. 5). *Ml*β-cat nuclear staining continues to be non-uniform until 4 hpf where nuclearization can be observed in cells at the oral pole, but nuclear staining was absent from the aboral pole.

**Fig. 5.**
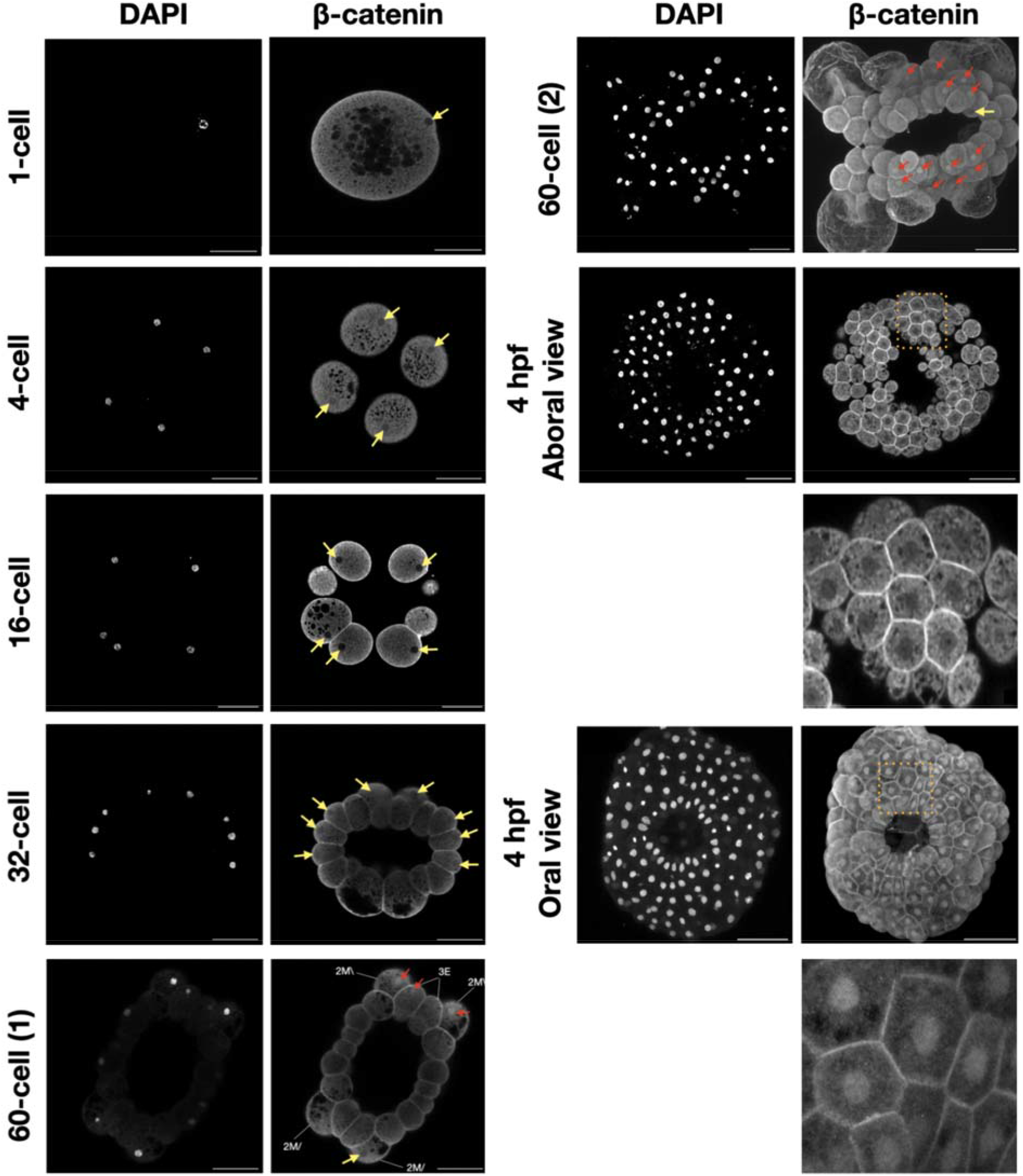
The localization of β-catenin in *M. leidyi* embryos. Immunostaining of *M. leidyi* embryos at different stages using affinity-purified β-cat antibodies. The left columns show nuclei stained by DAPI, the right columns show results of *Ml*β-cat antibody staining. Zoomed in insets (dotted boxes) for both the oral and aboral 4 hpf are included below the respective samples. The 1-cell, 4-cell, 16-cell, 32-cell, 60-cell (1), and 4 hpf aboral view are z-stack projected images. The 60-cell (2) and 4 hpf oral view are maximal projection images. The yellow arrows show absence of β-catenin nuclear translocation; while red arrows indicate β-catenin nuclear translocation. Scale bar, 50 µM.

## 4. Discussion

### 4.1 β-catenin robustly localizes to cell-cell interfaces in M. leidyi embryos

β-catenin involvement in the CCC is crucial for the adhesion between cells in every bilaterian animal where this process has been experimentally tested.^55^ Nevertheless, only a few studies focused on the roles of this complex in early branching lineages. Previous studies on the cnidarian *Nematostella vectensis* have shown that the CCC is involved in cell-cell adhesion and germ layer formation.^40,56^ Additionally, a functional CCC has been shown to be involved in cell-cell adhesion in sponges as supported by the co-immunoprecipitation of interacting CCC components, β-catenin localizing to cell-cell boundaries in *Ephydatia muelleri* and furthermore by *Yeast* 2-Hybrid screening in another sponge, *Oscarella carmela*.^57,58^

As for ctenophores, a prior study using polyclonal *Ml*β-cat antibodies generated against the first 10 amino acids of the protein was unable to capture evidence of the protein localized to the cell-cell interface in *M. leidyi*.^59^ Thus, it was concluded that the CCC is not conserved and hence does not play a role in cell-cell adhesion in ctenophores. However, there is a possibility that the short epitope used to generate the antibodies in the Salinas-Saavedra et al. 2019 study were being blocked by β-catenin binding partners at the cell surface, impeding the antibodies’ ability to bind and hindering the ability to observe this role. Unfortunately, the peptide antibody generated in the 2019 study appears to have lost its activity and we were not able to confirm its expression in embryos or Western blots (data not shown). Our results show that *Ml*β-catenin clearly localizes at cell-cell contacts during *M. leidyi* embryonic development and therefore likely has a cell-to-cell adhesion function in the earliest metazoan lineage (Fig. 5). Additionally, transcriptomic and genomic data showed that *M. leidyi* has a full set of CCC components and the amino acids essential for the interaction between β-catenin and ⍰-catenin and between β-catenin and cadherin are mostly conserved (Fig. 2).^46,47^ The conservation of almost all 12 Arm domains on *Ml*β-catenin leads us to predict that future functional studies will demonstrate that *Ml*β-catenin maintains its ability to interact with CCC partners in ctenophores.

However, it is yet unclear if a functional CCC exists in *M. leidyi* since the β-catenin binding site, essential for recruiting β-catenin at the membrane, is either cryptic or absent from *Mnemiopsis* cadherin sequences.^46,59^ An alternative hypothesis is that the CCC emerged from an interaction between β-catenin and a protocadherin instead of a classical cadherin. Indeed, it has been shown that a protocadherin can interact with β-catenin,^60–62^ but no study has shown that this interaction is necessary for cell-cell adhesion. A third option is cadherins are entirely unnecessary for cell-cell adhesion in ctenophores. An example of this was shown in the social amoeba *Dictyostelium discoideum* where in response to starving this organism undergoes the formation of a fruiting body, corresponding to a “multicellular” stage.^63^ Analysis of the *D. discoideum* genome indicates that ⍰-catenin and β-catenin genes are present while cadherins are absent. Knocking down the β*-cat* ortholog, called *Aardvark*, demonstrated that the catenins were essential for the establishment of this epithelium-like structure.^64^ Clarifying the interacting partners of *Ml*β-cat at the cell membrane will provide insights into the evolutionary origin of the CCC across metazoa.

### 4.2 β-catenin is nuclearized in cells around the blastopore in M. leidyi embryos

β-catenin can be found in three locations inside cells: in the cytoplasm, at the cell surface on the cytoplasmic side of the cell membrane, and in the nucleus. In each of these locations, it interacts with different proteins and maintains different roles. In order to fulfill its role as a cell signaling molecule, β-catenin must avoid being marked for degradation by the destruction complex in the cytoplasm and then translocate into the nucleus. In the vast majority of organisms where this process has been studied, this begins to occur in the early stages of development and facilitates transcription of genes involved in specifying cell fate.^27,28^ A prior study in *Mnemiopsis leidyi* tracked the spatiotemporal expression of the *Ml*β-catenin protein during development and found nuclearization beginning at the zygote stage.^59^ Furthermore, this study reported that there was continued nuclear expression of *Ml*β-cat in macromeres at the animal half in the endomesodermal progenitors through 11 hpf, arguing that this is substantial evidence to conclude that *Ml*β-cat has a role in specifying endomesoderm in early *M. leidyi* embryogenesis. When trying to replicate these results using the affinity-purified antibodies we generated to a *Ml*β-catenin polypeptide we were unable to find clear and consistent evidence of nuclear *Ml*β-cat in embryos prior to the 60-cell stage. From the 60-cell to 3 hpf we find nuclear *Ml*β-cat to be patchy and without a clear pattern. This is unexpected due to the highly stereotyped cleavage program and precocious specification of cell fate seen in ctenophores.^38^ Early stage ctenophore embryos develop at an accelerated pace making it difficult to capture specific stages for fixation and staining. This rapid cell cycle could also explain why visualizing nuclear β-catenin in these early embryos is difficult since the nuclear envelope is being disassembled and reassembled to accommodate mitotic processes. Improving methods for fixing may make obtaining spatial-temporal protein expression data more consistent in these early stages.

Scanning confocal analysis of *Ml*β-cat antibody stained *Mnemiopsis* embryos at around 4 hpf started to show a clear pattern of nuclear staining restricted blastomeres at the oral pole, while cells at the aboral pole did not show any nuclear staining. This is evidence that *Ml*β-cat may maintain its role as a co-activator of transcription, but does not align with the previously described expression pattern. The fact that this pattern is restricted to one pole does indicate the possibility for differential gene expression or stabilization, although patchy nuclear translocation during early development and a restriction to the ectodermal cells at the oral pole leaves us with questions about this protein’s functional role in cell type specification. Future functional experiments are required to determine if β-catenin maintains its role as a transcriptional co-activator in the Wnt/β-catenin pathway in ctenophores.

Furthermore, transportation of β-catenin into the nucleus is not well understood in all metazoans. There are several proposed mechanisms for transport including direct nuclear pore complex interaction, various piggyback or chaperone candidates, and post-translational modification.^65^ With their ancestral molecular repertoire and favorable optical properties, ctenophore embryos might be a powerful model to investigate the mechanisms behind β-catenin nuclearization.

### 4.3 Developing tools to study ctenophores

The combination of Western blot analysis and immunohistochemistry results gives us high confidence that the newly generated affinity-purified *Ml*β-cat antibody specifically targets β-catenin without any detectable non-specific interactions. As such, this antibody can now be used as a tool to ask follow-up questions about the functional roles of β-catenin in the earliest metazoan branch. The evolution and development community has come to recognize that ctenophores sit at a crucial point for understanding the early evolutionary history of metazoans and has been working to develop experimental methods and resources like assembled reference genomes, transcriptomes, morpholino oligonucleotides, and CRISPR-Cas9.^66–70^ Hopefully with this expanding toolkit we can begin to answer questions about transition from unicellularity to multicellularity in the animal lineage as well as better understand the molecular components the earliest metazoans maintain that are necessary for cell-cell adhesion and cell fate specification. As we develop new tools and techniques to study ctenophores, we will not only better understand how these animals operate on a molecular level, but also gain insights into how conserved mechanisms have evolved over time.

## Acknowledgements

This work was partially supported by NASA (80NSSC18K1067) to AHW and MQM and NSF (1755364) to MQM.

